# Screening for residues in Atg11, a central organizer of selective autophagy in yeast, important for binding with Atg9

**DOI:** 10.1101/2025.11.14.688473

**Authors:** Chimi Dolker Sherpa, Patricia Woghiren, Erin Leonello, Mina Abdel-Khalek, Benjamin J. Schuessler, Josh V. Vermaas, Steven K. Backues

## Abstract

The yeast protein Atg11, whose structure is unknown, is a central organizer of autophagosome formation that recruits Atg9 during selective autophagy. Although the residues in Atg9 responsible for this interaction are known, those in Atg11 are not. In an attempt to discover the binding site of Atg9 on Atg11, we screened a number of mutants within amino acid residues 455-627 of Atg11, guided in part by an AlphaFold2-generated model of the Atg11 dimer. However, we were not able to identify specific residues essential for the interaction with Atg9, suggesting that the binding region may lie elsewhere on Atg11.

## Description

Macroautophagy (hereafter referred to simply as autophagy) is a critical stress-response pathway conserved in all eukaryotes. Bulk autophagy, triggered by starvation, semi-randomly degrades materials from the cytoplasm in order to reuse the nutrients for cellular survival. In contrast, selective autophagy is not primarily regulated by a need for nutrients but targets specific cytoplasmic cargo to the lysosome (in animals) or vacuole (in other eukaryotes) for degradation (Xie & Klionsky, 2007). Selective autophagy contributes to the health of the cell by breaking down intracellular pathogens, protein aggregates, and malfunctioning organelles. In humans, it is particularly important for long-lived cells such as neurons and helps prevent neurodegenerative diseases such as Parkinson’s and Alzheimer’s (Choi et al., 2013; Lu et al., 2022). For this reason, there is great interest in understanding the proteins that control selective autophagy and how they interact. Much of this research has focused on autophagy in the model eukaryote *Saccharomyces cerevisiae*, baker’s yeast.

Selective autophagy in yeast begins when cargo receptor proteins are recruited to a specific autophagic cargo. These receptors bind to Atg11, the central organizer of the autophagy initiation complex during selective autophagy (Yorimitsu & Klionsky, 2005; Zientara-Rytter & Subramani, 2020). This then arranges the proteins that start the process of building the autophagosome - the double-membrane vesicle that will envelop the cargo and deliver it to the vacuole for degradation.

The structure of Atg11 is unknown, as is the manner by which it recruits and organizes the autophagic machinery. It is predicted to be a dimer with many coiled-coil domains (Yorimitsu & Klionsky, 2005). Biophysical data has verified it to be a rod-shaped parallel dimer (Suzuki & Noda, 2018), and a crystal structure of a portion of coiled-coil domain 3 shows it to be a flexible coil (Margolis et al., 2020). Atg11 binds to cargo receptors via its C-terminal claw domain, the structure of which has been solved for its human homolog FIP200 (Turco et al., 2019). A cryo-EM structure the N-terminal half of the FIP200 protein is also available (Chen et al., 2025); however, the N-terminal region of FIP200 is not homologous to that of Atg11. This region binds to core autophagic machinery such as the kinase Atg1 and the scramblase Atg9 (He et al., 2006; Meyer et al., 2022; Yorimitsu & Klionsky, 2005). Atg9 is a trimeric transmembrane protein that is found in populations of small vesicles, and Atg11 binding to Atg9 recruits these vesicles and causes their fusion to form the phagophore (Backues et al., 2015; Yamamoto et al., 2012). Atg9 equilibrates lipids delivered from the ER to both bilayers of this membrane; as such, it is critical for the formation of the autophagosome (Guardia et al., 2020; Orii et al., 2021; Sawa-Makarska et al., 2020).

Atg9 has been shown to interact with Atg11 via two PLF motifs in its N-terminal unstructured region: PLF^163-165^ and PLF^187-189^ (Coudevylle et al., 2022). A neighboring histidine, H192, is also involved (He et al., 2006), and mutation of these residues blocks autophagy via preventing recruitment of Atg9 to the site of autophagosome formation (Coudevylle et al., 2022; He et al., 2006). However, it is unknown which residues in Atg11 interact with Atg9.

Previous studies have shown that amino acid residues (a.a.) 1-627 of Atg11 are sufficient for binding to Atg9 (He et al., 2006) while a.a. 1-454 are not (Hollenstein et al., 2021), suggesting that the Atg9 binding site might be between a.a. 455-627. Our lab previously investigated part of this region: a.a. 536-576, termed coiled-coil domain 2 (CC2), which had initially been implicated as a likely site for the binding of both Atg1 and Atg9 as well as Atg11 dimerization (Yorimitsu & Klionsky, 2005). However, our systematic mutagenesis data suggested that CC2 is more likely to be involved in overall folding and stability of the N-terminus of Atg11 (Meyer et al., 2022). Therefore, we focused on a.a. 455-535, as that region had not been previously investigated (Figure 1A).

**Figure 1.**
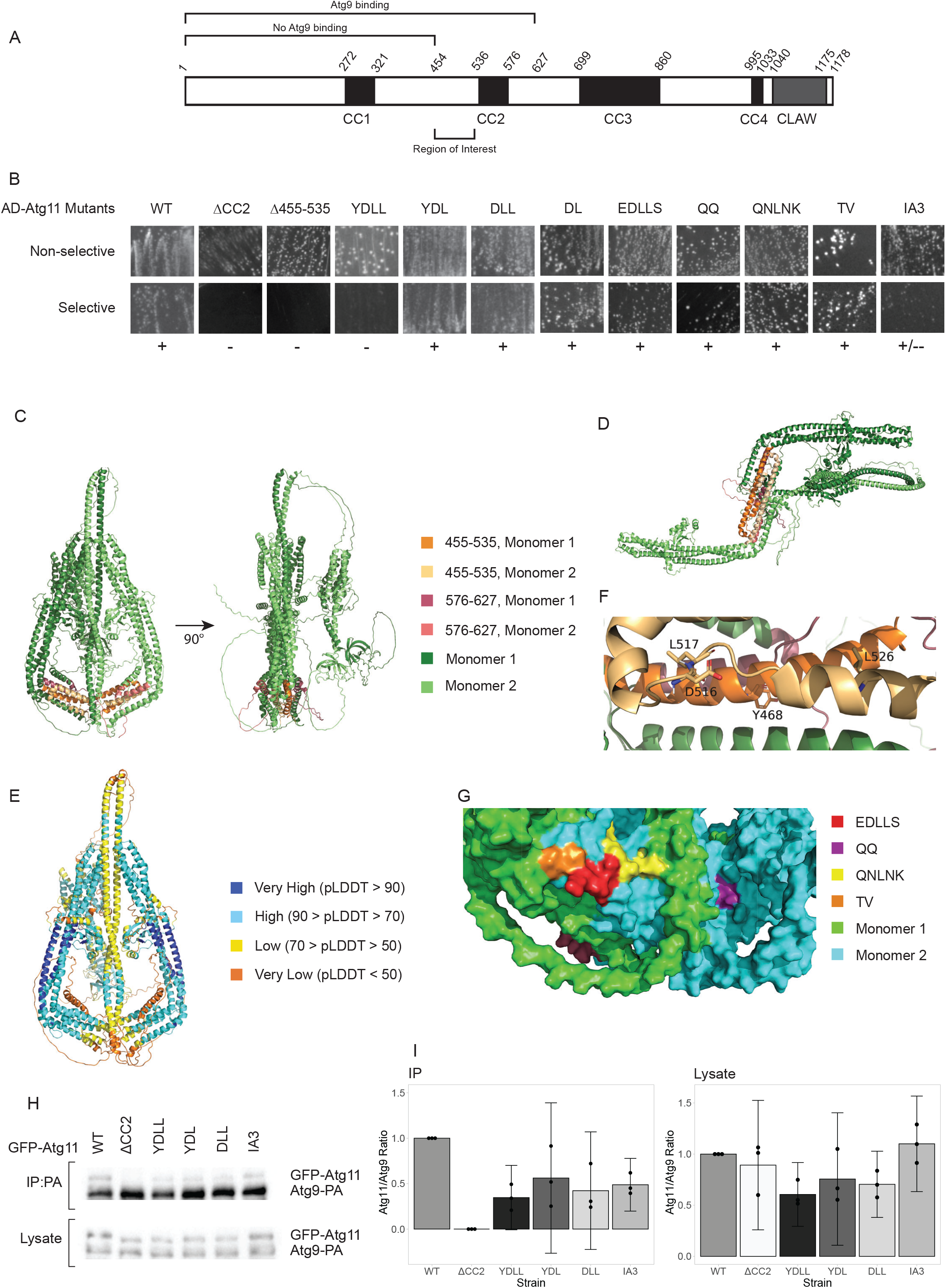
Mutations in the 455-535 and 576-627 regions of Atg11 do not cause full loss of interaction with Atg9: **A**Schematic representation of Atg11 showing the coiled-coil domains (black boxes) and the CLAW domain (grey box), with a.a. shown by italicized numbers. Fragments previously reported to bind or not bind with Atg9 are indicated. **(B)** Yeast 2 hybrid assay of BD-Atg9 with the indicated AD-Atg11 mutants. Non-selective media is SMD -ura -leu; selective media is SMD -ura -leu -his +1.5 mM 3-AT. On selective media, (+) indicates growth and (-) indicates no growth across multiple replicates of the assay. **(C)** Front and side view of the flask-shaped Atg11 dimer and **(D)** front view of the Z-shaped Atg11 dimer predicted with AlphaFold2 Multimer. **(E)** pLDDT confidence map of the flask-shaped Atg11 structure. **(F)** Close-up of flask-shaped dimer showing predicted locations of YDLL residues. **(G)** Surface rendering of flask-shaped model highlighting predicted surface exposed residues in the 455-535 region. **(H)** CoIP of various overexpressed GFP-Atg11 mutants by overexpressed PA-Atg9. Blots were probed with anti-GFP, which also detects the PA tag. **(I)** Quantitative analysis of CoIP results. The Atg11 to Atg9 band intensity ratio was normalized to the values from WT for each replicate. For the CoIP graph, the GFP-Atg11 band intensity of the ΔCC2 samples, which are known to show no binding (Meyer et al., 2022), were used as the background and subtracted from the GFP-Atg11 band intensity of the other samples before normalizing. Error bars are 95% confidence intervals.

Yeast-2-Hybrid (Y2H) results showed that atg11Δ455-535 lost interaction with Atg9, just like atg11ΔCC2 (Figure 1B). We then examined sequence of the 455-535 region across various yeast species, looking for residues that were conserved in the family *Saccharomycetacea*, where the Atg11-binding region of Atg9 was conserved, but not conserved in other species from order *Saccharomycetales* where the Atg11-binding region of Atg9 was not conserved. This analysis yielded four residues: Y468, D516, L517 and L526. A quadruple mutant of all of those residues to alanine (YDLL) mostly lost interaction with Atg9 by yeast-2-hybrid. However, subsets of 2 or 3 of these mutations (YDL, DLL, or DL) did not lose interaction, suggesting that none of these was a critical residue (Figure 1B).

We used AlphaFold2 Multimer to predict the structure of the Atg11 dimer. The top-ranked model was a “flask-shaped” parallel dimer (Figure 1C). However, some other models predicted a Z-shaped dimer (Figure 1D). Both models showed extensive monomer-monomer interactions in CC2, CC3 and CC4, consistent with previous reports that both the N- and C-termini of Atg11 could dimerize (Suzuki & Noda, 2018). They both suggested that CC1 and CC2 formed an L-shaped helical bundle, with CC3 and the C-terminus connected by disordered linkers; however, the placement of CC3 and the C-terminus varied between these models - in the flask-shaped, they were neatly packed against each other, while in the Z-shaped dimer, they were placed at random angles, not interacting with CC1, CC2, or each other. The pLDDT for both models was relatively low, particularly at a critical interface at the bottom of the flask-shape, raising the possibility that Atg11 can take on multiple conformations (Figure 1E).

Y468 and L526 were placed on the interior of the Atg11 dimer in both AlphaFold models, which would not allow a direct role in Atg9 binding, while D516 and L517 were partially surface exposed (Figure 1F). We mutated residues E515-S519 to alanine (EDLLS) to target that entire surface, but this mutant did not lose interaction with Atg9 (Figure 1B). We also generated three additional mutants, QQ, QNLNK, and TV, all of which targeted sets of residues within the 455-535 region that the AlphaFold models predicted to be surface exposed (Figure 1G). However, none of these mutants lost interaction, either (Figure 1B).

The region downstream of CC2 was primarily disordered in our AlphaFold models. To determine if this region was important for Atg9 binding, we replaced residues 576-627 of Atg11 with residues 2-53 of an intrinsically unstructured protein, PAI3. The Atg11_IA3 mutant showed a partial loss of interaction with Atg9 by yeast-2-hybrid (Figure 1B).

Finally, we confirmed our yeast-2-hybrid results by performing co-immunprecipitation (CoIP) assays between Atg9-PA and selected GFP-Atg11 mutants. These showed a partial loss of interaction between Atg9 and all of the Atg11 mutants tested - YDLL, YDL, DLL and IA3. In addition, there was a reduction of the levels of the YDLL, YDL and DLL mutants in the lysates, suggesting that these mutations might be detrimental to the overall stability of Atg11 (Figure 1H,I).

Overall, none of the residues or regions we mutated emerged as strong candidates for an Atg9 binding site. The YDLL mutant showed a loss of interaction only when all four residues were mutated, and CoIP results showed that this was only a partial loss of interaction. The fact that these residues are predicted to be in different locations on the folded protein and two of them are predicted to be interior-facing, along with the lower expression observed for this mutant, suggests that they have a primary effect on Atg11 folding and only a secondary effect on Atg9 binding. Similarly, the fact that replacement of 52 residues with those of an unrelated protein led to only a partial loss in binding suggests that the a.a. 576-627 is also not the binding site for Atg9. Interestingly, a recent study showed that an overlapping region, a.a. 612-646, has membrane-binding properties, supporting a different role for this part of Atg11 (Andhare et al., 2025).

We have not exhaustively mutated a.a. 455-627, so we can not rule out the possibility that Atg9 binds in this region. However, we have mutated the most likely candidate residues without finding any specifically responsible for Atg9 binding, suggesting that the core Atg9 binding region may lie elsewhere in the Atg11 N-terminus, while a.a. 455-627 have an indirect effect. The AlphaFold multimer model predicts a.a. 454-591 to be a dimerization interface for Atg11; therefore, we speculate that the removal of a.a. 455-627 from the N-terminal construct may disrupt its proper folding and dimerization, leading indirectly to a loss of Atg9 binding.

## Methods

### Strain and plasmid construction

The pGAD-atg11 mutant plasmids were ordered from GenScript. The pRS416-Cu-GFP-atg11 mutant plasmids (YDLL, YDL, and DLL) were made using InFusion cloning. The vector pRS416-Cu-GFP-ATG11 was digested with EcoRI-HF (NEB), which cuts at two endogenous EcoRI sites in Atg11. The insert was amplified by PCR using Phusion polymerase (NEB) from the respective pGAD-Atg11 mutant with primers agaagaagaagaattcaatagtcaag and atttgctatagaattctcattttcatc, ordered from IDT. The vector and insert were combined by InFusion reaction (Clontech, Takara Bio USA Inc.) and transformed into E. coli DH5α (Stellar Competent Cells - HST08 strain) and plated on LB carbenicillin plates for colony growth. pRS416-Cu-GFP-Atg11^IA3^ was generated by similar methods, except the digest was performed with AvrII and HindIII and the primers used were cagatggaaatcctaggtatta and caacttcataaagcttattgtc.

Plasmids were purified using Purelink plasmid miniprep kits (ThermoFisher), verified by Sanger sequencing (Eton Biosciences) and/or whole plasmid sequencing (Plasmidsaurus) and introduced into yeast cells using standard transformation procedures (Gietz & Schiestl, 2007).

### Yeast-2-hybrid

Yeast multiple-knockout cells (lacking 25 *ATG* genes to minimize indirect or interfering interactions; YCY149) expressing BD-ATG9 and AD-ATG11 or mutants thereof were patched on both selective (SMD -ura -leu -his w/ 1.5 mM 3-amino-triazol) and nonselective (SMD -ura -leu) plates and grown for 4–5 days at 30°C before imaging. Results shown are representative of multiple replicates from multiple independent transformants.

### Co-Immunoprecipitation and Western Blotting

Co-immunoprecipitation was carried out exactly as described previously (Meyer et al., 2022) except that all washes were performed in the 4℃ cold room to help prevent breakdown. Precast Bio-Rad 4-15% gradient SDS-PAGE gels and GoldBio BLUEstain2 protein ladder were used. The electrophoresis was run for ∼3 hours, starting at 80 V and increasing to 120 V after initial band separation.

### Structural Modeling

To develop structure-based hypotheses for potential interaction sites between Atg9 and Atg11, AlphaFold-Multimer version 2.3.2 (Evans et al., 2022) was used to generate 25 potential interaction complexes. These complexes were optimized using the FoldX4 “repair” function (Delgado et al., 2019). Visualization was performed in PyMol 3 (Schrödinger, LLC, 2015).

**Table 1:** Strains and Plasmids used in this study.

| ***Saccharomyces cerevisiae* strains used in this study** |  |  |
| --- | --- | --- |
| **Name** | **Genotype** | **Source** |
| atg11Δ | MATα *his3Δ200 leu2-3,112 lys2-801 suc2-Δ9 trp1Δ901 ura3-52 atg11Δ::KanMX* | Meyer et al., 2022 |
| YCY149 | MATα *his3-Δ200 leu2-3,112 lys2-801 trp1-Δ901 suc2-Δ9 ura3-52 atg1Δ, 2Δ, 3Δ, 4Δ, 5Δ, 6Δ, 7Δ, 8Δ, 9Δ, 10Δ, 11Δ, 12Δ, 13Δ, 14Δ, 16Δ, 17Δ, 18Δ, 19Δ, 20Δ, 21Δ, 23Δ, 24Δ, 27Δ, 29Δ gal4Δ gal80Δ GAL1-HIS3 GAL2-ADE2 met2::GAL7-lacZ atg31Δ::KanMX* | Cao et al., 2009 |
| **Plasmids used in this study** |  |  |
| **Name** | **Description** | **Source** |
| pGBDU-ATG9 | BD vector with full length ATG9 | He et al., 2006 |
| pGAD-ATG11 | AD vector with full length WT ATG11 | Meyer et al., 2022 |
| pGAD-atg11ΔCC2 | AD vector with atg11Δ<sup>536-576</sup> | Meyer et al., 2022 |
| pGAD-atg11Δ455-535 | AD vector with atg11Δ<sup>455-535</sup> | This study |
| pGAD-atg11<sup>YDLL</sup> | AD vector with atg11<sup>Y468A, D516A, L517A, L526A</sup> | This study |
| pGAD-atg11<sup>YDL</sup> | AD vector with atg11<sup>Y468A, D516A, L517A</sup> | This study |
| pGAD-atg11<sup>DLL</sup> | AD vector with atg11<sup>D516A, L517A, L526A</sup> | This study |
| pGAD-atg11<sup>DL</sup> | AD vector with atg11<sup>D516A, L517A</sup> | This study |
| pGAD-atg11<sup>EDLLS</sup> | AD vector with atg11<sup>E515A, D516A, L517A, L518A, S519A</sup> | This study |
| pGAD-atg11<sup>QQ</sup> | AD vector with atg11<sup>Q475A, Q476A</sup> | This study |
| pGAD-atg11<sup>QNLNK</sup> | AD vector with atg11<sup>Q493A, N494A, L496A, N500A, K501A</sup> | This study |
| pGAD-atg11<sup>TV</sup> | AD vector with atg11<sup>T511A, V512A</sup> | This study |
| pGAD-atg11<sup>IA3</sup> | AD vector with atg11 where residues 576-627 were replaced with residues 2-53 of *scPAI3* (aka IA3). | This study |
| pRS414-Cu-ATG9-PA | ATG9 tagged with Protein A, driven by the *CUP1* promoter | Meyer et al., 2022 |
| pRS416-Cu-GFP-ATG11 | GFP-ATG11 WT driven by the *CUP1* promoter | Kim et al., 2001 |
| pRS416-Cu-GFP-Atg11ΔCC2 | GFP-atg11$\Delta^{536-576}$ driven by the *CUP1* promoter | Meyer et al., 2022 |
| pRS416-Cu-GFP-Atg11<sup>YDLL</sup> | GFP-atg11<sup>Y468A, D516A, L517A, L526A</sup> driven by the *CUP1* promoter | This study |
| pRS416-Cu-GFP-Atg11<sup>YDL</sup> | GFP-atg11<sup>Y468A, D516A, L517A,</sup> driven by the *CUP1* promoter | This study |
| pRS416-Cu-GFP-Atg11<sup>DLL</sup> | GFP-atg11<sup>D516A, L517A, L526A</sup> driven by the *CUP1* promoter | This study |
| pRS416-Cu-GFP-Atg11<sup>IA3</sup> | GFP-atg11 with residues 576-627 replaced with residues 2-53 of *scPAI3* (aka IA3). Driven by the *CUP1* promoter. | This study |
| **Antibodies used in this study** |  |  |
| **Name** | **Identifier (Source)** | **Concentration** |
| “Living Colors” mouse monoclonal anti-GFP (JL-8) | Cat# 632381 (Clontech / Takara) | 1:500 |
| Peroxidase AffiniPure® Goat Anti-Mouse IgG | Cat# 115-035-003 (Jackson ImmunoResearch) | 1:5000 |

## Acknowledgements

This research was supported in part by the National Science Foundation under RUI grant 2243163 to S.K.B and the National Institute of General Medical Sciences of the National Institutes of Health under award number R35GM155317 to J.V.V.

